# S-adenosylmethionine supply controls DNA methylation turnover in synovial sarcoma

**DOI:** 10.64898/2026.11.09.765438

**Authors:** Yanxiang Li, Md Salman Shakil, Matthew E. Johnson, M. Elena Díaz-Rubio, Katarzyna Kosicka-Noworzyń, Leonid Kagan, Luigi Brunetti, Xiaoyang Su, Matthew J. McBride

## Abstract

**Background:** Methionine is the nutrient source of the metabolite S-adenosylmethionine (SAM) used to methylate chromatin for regulation of gene expression. Disruptions to this metabolic supply, including nutritional deficiency of certain B vitamins, alter chromatin methylation and have been implicated in tumor growth.

**Objective:** To define how cellular SAM supply controls DNA methylation dynamics and the disruption of oncogenic gene expression in synovial sarcoma.

**Methods:** Cell line models of synovial sarcoma were treated with the DNA methyltransferase inhibitor azacitidine or the small-molecule MAT2A inhibitor AG-270, which blocks SAM synthesis from methionine. Intracellular SAM concentrations were quantified by liquid chromatography-mass spectrometry. Global DNA methylation rates were measured by stable isotope tracing with [^2^H_3_-methyl]-methionine. Genomic sites of altered DNA methylation were identified by EM-seq and the associated transcriptional changes measured by RNA-seq.

**Results:** Inhibition of SAM synthesis reduced intracellular SAM concentration from 94 µM to 5.6 µM (p value < 0.0001) at 24 hours. This was associated with a 19% decrease (p value < 0.0001) in the global 5-methyldeoxycytosine turnover rate, while total DNA methylation levels were maintained. In contrast to the global DNA hypomethylation induced by azacitidine, SAM depletion resulted in a redistribution of DNA methylation with loss at 252 and gain at 206 genomic sites, and the transcriptional response to SAM depletion was not associated with changes in promoter methylation at the altered genes. Both azacitidine and SAM depletion activated mesenchymal genes, while suppressing cell cycle progression.

**Conclusions:** Metabolic supply of SAM is a major regulator of DNA methylation turnover providing a mechanism by which nutritional or genetic disruptions to SAM metabolism can alter chromatin methylation and gene expression.

## INTRODUCTION

Methionine is an essential amino acid required for protein synthesis and is connected by B-vitamin nutrients to the folate cycle to control availability of one-carbon units(1–3). Methionine directly contributes to cellular methylation reactions through its conversion into S-adenosylmethionine (SAM) by methionine adenosyltransferase (MAT) enzymes(4). SAM serves as the universal methyl-group donor for DNA, RNA, and histone tail methylation(5, 6). Through these methylation-dependent processes, SAM supply alters chromatin structure and accessibility to regulate gene expression(7, 8). Consequently, nutrient availability and/or one-carbon metabolism enzyme activity that alters SAM supply can have a direct influence on chromatin methylation(9, 10).

Restricting exogenous or dietary methionine availability in cell lines, rodent models and humans directly results in reduced methionine supply to disrupt SAM levels(7, 11–15). Folate (vitamin B9) and cobalamin (vitamin B12) deficiency prevent remethylation of homocysteine for methionine generation leading to reduced SAM pools, particularly in the liver and brain(16–19). Supplementation of these B vitamins rescues this methylation-deficiency(20, 21). Similarly, mutations to one-carbon metabolism genes also disrupt SAM supply. Reduced function mutations to *MTHFR* (Methylenetrahydrofolate Reductase) and *MTR* (Methionine Synthase) directly slow transfer of the one-carbon unit from the folate cycle for methionine generation(22, 23). Mutations to *AHCY* (S-adenosylhomocysteine hydrolase) or *CBS* (cystathionine β-synthase) cause an accumulation of the methionine cycle intermediates S-adenosylhomocysteine (SAH) and homocysteine impairing metabolic utilization of SAM(24–26).

Chromatin structure, gene expression and resulting cellular identity are linked to these nutritional and genetic driven alterations in metabolic SAM supply through changes in methylation. Epigenetic methylation has distinct regulatory roles that are dependent on the specific macromolecule and position of the methylation(27). Methylation of cytosine residues of DNA in gene promoters reduces accessibility to drive gene silencing, while emerging work shows that DNA methylation in gene bodies contributes to gene activation(16, 28). Folate deficiency has generally been associated with global DNA hypomethylation, with specific sites of hypermethylation, yet the relationship between DNA methylation and folate status is appreciated to be complex(29, 30). Similarly, people with MTHFR mutations in combination with a nutritional folate-deficiency are associated with reduced global DNA methylation levels(31–34). Beyond DNA, SAM metabolism influences RNA and histone methylation. Folic acid supplementation in mice has led to elevated hepatic levels of specific methylated mRNA transcripts(35). Histone methylation is likely the most sensitive to alterations in SAM supply(7). Methylation of lysine and arginine residues on histone protein tails of nucleosomes alter recruitment of transcription factors and other transcriptional protein machinery to chromatin contributing to both gene activation and suppression(36, 37). Methionine restriction in both cell lines and mouse models has shown a reduction in histone methylation, particularly trimethylation of lysine 4 on histone 3 (H3K4me3)(7).

A major contributor to the complex relationship between SAM metabolism, chromatin methylation and transcriptional output is that methylation is not a static modification, but a dynamic mark that undergoes continuous turnover(38, 39). The steady-state level of methylation is determined by the balance between methylation and demethylation rates, which can vary at distinct sites across the genome(40). To maintain methylation levels over time, these opposing processes are balanced to tune transcriptional output. Work in mouse embryonic stem cells shows that genes with high transcriptional activity are associated with elevated rates of DNA methylation turnover driven by high demethylation rates(41, 42). Many approaches for measuring methylation turnover commonly rely on genetic or pharmacological perturbation of methylation enzymes followed by monitoring the changes in DNA methylation levels over time. Here, we adopt a stable isotope tracer-based approach to directly quantify DNA methylation turnover fluxes under conditions of reduced SAM availability.

Numerous studies show that restriction of exogenous methionine supply reduces tumor growth and can enhance cytotoxic chemotherapy and radiation(1, 15). Recently, it has been discovered that approximately 15% of all cancers harbor a deletion at chromosome 9p21 resulting in enhanced sensitivity to tumor SAM depletion(43). This has motivated multiple Phase I/II clinical trials to evaluate the tolerability and efficacy of small molecule inhibitors targeting MAT2A, the major source of SAM synthesis from methionine outside the liver(44). However, the effects of MAT2A inhibition on chromatin methylation are undefined and there is evidence in gliomas and leukemia that targeting MAT2A can have therapeutic benefit even in cancers that lack of this 9p21 chromosomal alteration(45, 46). Consequently, here, we use the MAT2A inhibitor AG-270 to disrupt SAM supply and quantify DNA methylation rates in the cancer synovial sarcoma.

Synovial sarcoma is an aggressive soft tissue sarcoma defined by a hallmark chromosomal translocation that results in formation of the SS18-SSX fusion oncoprotein(47). This SS18-SSX fusion incorporates into SWI/SNF (BAF) nucleosome remodeling complexes to activate bivalent genes, prevent mesenchymal differentiation, and drive oncogenesis in a rare fibroblastic mesenchymal stem cell population(48–51). This suggests that the oncogenic activity of SS18-SSX is highly dependent on a specific epigenetic state permissive to transformation. Consistent with this, DNA hypomethylation by azacitidine treatment disrupts SS18-SSX localization on chromatin to induce mesenchymal gene activation and arrest tumor growth(52). Here, we report that SAM depletion by pharmacological targeting of MAT2A reduces DNA methylation turnover fluxes and this causes a balanced redistribution of DNA methylation genome-wide, induces differentiation, and blocks synovial sarcoma cell growth.

## METHODS

### Cell Culture

HSSYII, established from a monophasic synovial sarcoma tumor expressing *SS18-SSX1*, was obtained from RIKEN Bio Resource Research Center (Tsukuba, Japan) and SYO1, established from a biphasic synovial sarcoma tumor expressing *SS18-SSX2*, was a gift from Cigall Kadoch (Dana Farber Cancer Institute)(53, 54). HSSYII and SYO1 cells were cultured in DMEM (Gibco, 11965118) with 10% fetal bovine serum (FBS) (Gibco) and 1% penicillin/streptomycin (Gibco). Cells were maintained at 37°C in a humidified atmosphere with 5% CO2 and screened for mycoplasma contamination every 3 months.

### Drug Treatments and Proliferation Assay

HSSYII or SYO1 cells were plated in 6 well plates at a density of 100,000 cells per well. After 24 hours, cells were treated at a final concentration in the culture media with Vehicle (0.1% DMSO), 6 µM AG-270 (MedChemExpress, HY-138630), 3 µM AG-270, or 1 µM azacitidine (MedChemExpress, HY-10586) for 72 hours or 144 hours, with drug-containing culture media refreshed every 72 hours. Cells were then fixed with 10% trichloroacetic acid (TCA) at 4°C for 30 minutes and gently washed twice with distilled water. The plates were dried, stained with Sulforhodamine B (SRB) dye, incubated at room temperature for 30 minutes, and washed twice with 1% acetic acid. The plates were dried, SRB dye dissolved in Tris base pH 10.5 solution and 565 nm absorbance measured by microplate reader.

### SAM Quantification by LC-MS/MS

Cells in 6 well plates were washed once with PBS, and intracellular metabolites extracted in 600 µL of cold methanol:water (80:20) with 1 mM N-ethylmaleimide (NEM). 25 µM of ^13^C-methyl labeled SAM (MedChemExpress, HY-B0617S2) internal standard was spiked into each extraction at 1 µL per µL of pack cell volume. The cell extracts were centrifuged at 4°C for 30 min at 16,000 × g and supernatant transferred to LC-MS vial. The metabolite extract was then analyzed by hydrophilic interaction chromatography (HILIC) coupled to a QTRAP 6500+ (Sciex) mass spectrometer with electrospray ionization. The LC separation was by a XBridge Premier BEH Amide VanGuard FIT column (150 mm × 2.1 mm, 2.5 μm particle size, Waters, 186009933) with a gradient of solvent A (95%:5% H2O:acetonitrile with 20 mM acetic acid, 40 mM ammonium hydroxide, pH 9.4) and solvent B (20%:80% H2O:acetonitrile with 20 mM acetic acid, 40 mM ammonium hydroxide, pH 9.4). The gradient was 0 min, 100% B; 3 min, 100% B; 3.2 min, 90% B; 6.2 min, 90% B; 6.5 min, 80% B; 10.5 min, 80% B; 10.7 min, 70% B; 13.5 min, 70% B; 13.7 min, 45% B; 16 min, 45% B; 16.5 min, 100% B; and 22 min, 100% B. The injection volume was 5 µL, flow rate was 300 μl per min, column temperature was 25 °C, and the autosampler temperature was 4 °C. MS scans were obtained in positive mode ionization with corresponding parent (399 m/z) and daughter (250 m/z) ions detected at 10.5 min for SAM. Curtain gas (CUR) was at 30, collision gas (CAD) at Medium, ion spray voltage (IS) at 3500 Volts, temperature (TEM) at 450 °C, ion source gas 1 (GS1) at 60 and ion source gas 2 (GS2) at 60. Quantification of metabolites was performed using Analyst^®^ 1.6.3 software.

### Stable Isotope Tracing

Cells were cultured in 6 cm dishes or 6-well plate, and after 24 hours, the media was replaced with DMEM without methionine and choline chloride (USBiological Science, 38210000) supplemented with 30 µM choline chloride (Millipore Sigma), 200 µM [^2^H_3_-methyl]-methionine (Cambridge Isotope Laboratories, DLM-10774), and 10% dialyzed FBS (Gibco). All cells were cultured with isotope tracers for 24 h, 48 h, or 72 h followed by harvesting for metabolite or DNA extractions.

### DNA and RNA Extraction

Cells were collected by scraping into culturing media, transferring to tube, centrifuging for 3 minutes at 1,400xg and aspirating off the media from the cell pellet. The cells were washed with PBS and resuspended in 500μL of Zymo Research DNA/RNA shield buffer by pipetting up and down 10 times and stored at –80**°**C until DNA and RNA was isolated. DNA and total RNA was simultaneously isolated from the frozen cells following Zymo Research Quick-DNA/RNA Microprep Plus Kit (Cat#: D7005) protocol with no deviations from the kit protocol. After isolation, DNA and total RNA was stored at –80**°**C.

### DNA Digestion

1 µg of extracted DNA was digested with the nucleoside digestion mix (New England Biolabs, M0649S) at 37 °C for 1 hour. Digested DNA was then diluted 1:10 in 100% HPLC-grade methanol, centrifuged at 4 °C 16,000 × g for 10 min, and supernatant transferred to LC-MS vial.

### Nucleoside Analysis by LC-MS

Following DNA digestion, nucleosides were analyzed on an Oribitrap IQ-X mass spectrometer (Thermo) by electrospray ionization. Samples were separated by same LC HILIC column and method as previously described (see ***SAM Quantification by LC-MS/MS*** section). Injection volume was 5 µL and MS scans were obtained in positive ion mode with a resolution of 120,000 at m/z 200, automatic gain control target of 100%, and a m/z scan range of 72 to 1000. Metabolite data was annotated and analyzed using El-MAVEN with a mass accuracy window of 10 ppm for peak selection. Isotope natural abundance correction was performed using AccuCor(55).

### Transcriptomics Analysis (RNA-seq)

Total RNA underwent library generation and sequencing by NovoGene. Briefly, RNA-seq libraries were prepared using NEBNext Ultra II Directional RNA Library Prep Kit for Illumina and NEBNext Poly(A) mRNA Magnetic Isolation kit for poly A transcript isolation following standard protocols. The libraries were sequenced on a NovaSeq X Plus Series (PE150bp) to a depth of 70 million reads per sample. Adaptor sequences and any reads less than 10bp in length were removed. The sequence reads were aligned to UCSC human genome build hg38 by STAR aligner (version 2.7.5a). HOMER (Hypergeometric Optimization of Motif EnRichment) version 5.1 was used to generate raw read coverage and RPMK files for all transcripts. The raw read files were analyzed using DEBrowser v1.30.2 for differential expression analysis. Low count filtering was done based on mean when max value was less than 10 and batch effect normalization used RLE with correction method CombatSeq.

### Global DNA Methylation Profiling (EM-seq)

Genomic DNA underwent library generation and sequencing by NovoGene. Briefly, EM-seq libraries were prepared using NEBNext Enzymatic Methyl-seq v2 Kit following standard protocols. The libraries were sequenced on a NovaSeq X Plus Series (PE150bp) to a coverage depth of 30X human genome. Adaptor sequences were trimmed and any reads less than 10bp in length were removed. The sequenced reads were analyzed using Bismark version 0.24.2 for alignment of sequence reads to UCSC human genome build hg38 with bowtie 2 command options (-score_min L,0,-0.6). After alignment, deduplication was performed using Bismark’s deduplication tool. Bismark methylation extractor was used on the deduplicated alignment files to extract the methylated and unmethylated CpGs in the aligned sequence reads to generate Bismark coverage files. Then coverage files were analyzed using SeqMonk version 1.48.1 using UCSC human genome build hg38 for reference alignment genome(56). Differentially Methylated Regions (DMRs) were determined using SeqMonk’s read position probe generator requiring ≥10 CpG sites within a 500bp region. DMRs were further filtered for statistical significance using logistic regression on the per-CpG read positions with regions considered significant at p ≤ 0.05 and a normalized methylation difference of ≥20 percentage points (hypermethylation) or ≤-20 percentage points (hypomethylation) between conditions.

### Statistical analysis

All data are shown as mean ± SEM. For comparisons of two groups, statistical significance was calculated using a two-tailed, unpaired Student’s t test. For comparison of multiple treatment groups, one-way ANOVA with Dunnett’s multiple comparison test was carried out. All statistical analysis was performed using GraphPad Prism (version 10.5.0).

## RESULTS

### MAT2A inhibition depletes SAM levels

In non-hepatic tissue, SAM is synthesized from methionine by MAT2A and then this metabolite is used as a substrate by methyltransferases to methylate cytosine residues of DNA (Fig. 1A). We found that treating the HSSYII and SYO1 synovial sarcoma cell line models with small molecule inhibitors of MAT2A (AG-270) and DNA methyltransferases (azacitidine) reduced cellular proliferation (Fig. 1B-C, Fig. S1A-D). To understand how these treatments alter cellular SAM supply, we performed liquid chromatography-mass spectrometry (LC-MS) to quantify SAM pools. While SAM levels were modestly reduced by azacitidine treatment, targeting MAT2A dramatically depleted SAM supplies (Fig. S1E). By absolute quantification, we found that intercellular SAM concentrations are 94 µM under standard nutrient-rich culture conditions and following MAT2A inhibition are reduced to 5.6 µM within 24 hours (Fig. 1D, Fig. S1F-H). To consider how this reduced supply of SAM could impact the kinetics of DNA methylating enzymes, we compared these measured concentrations to reported K_m_ values of DNMT1 and DNMT3A/B for the substrate SAM (Fig. 1E)(57). While SAM levels of cells grown under standard culture conditions are in far excess, reducing SAM levels to low micromolar is near the range of reported K_m_ values and could be expected to impact DNA methylation rates.

**Figure 1.**
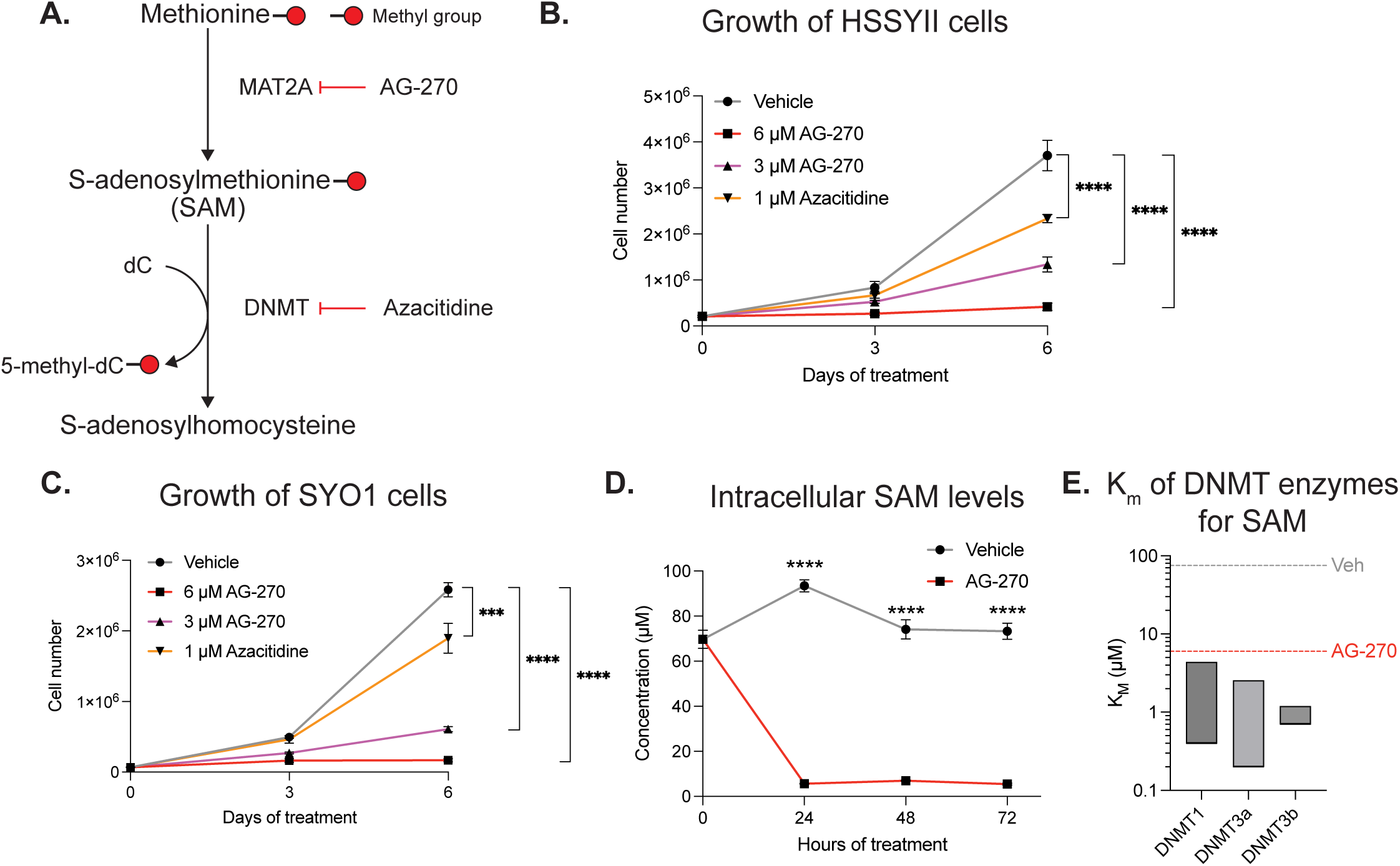
Inhibition of MAT2A reduces intracellular SAM levels to block proliferation of synovial sarcoma cells. (A) Pathway schematic of methionine cycle showing targets of small molecule inhibitors AG-270 for MAT2A and azacitidine for DNMT. (B) HSSYII and (C) SYO1 synovial sarcoma cell line growth following vehicle, 3 or 6 µM AG-270, or 1 µM azacitidine treatment for 3 or 6 days (n=3). (D) Intracellular SAM concentrations quantified by LC-MS/MS in HSSYII after vehicle or 6 µM AG-270 treatment for 24, 48, and 72 hours (n=4). (E) K_m_ values of DNA methyltransferases DNMT1, DNMT3a, and DNMT3b for SAM compared to measured SAM pools from (D). All data are reported as mean ± SD, p value by one-way ANOVA with Dunnett’s multiple comparison test and replicates (n) indicated. * denotes p < 0.05, ** denotes p < 0.01, *** denotes p < 0.001, **** denotes p < 0.0001.

### Decreased SAM supply slows DNA methylation turnover rates

To quantify the impact of reduced SAM supply on DNA methylation levels and rates, we paired LC-MS analysis with stable isotope tracing (Fig. 2A). We cultured HSSYII and SYO1 cells under standard culture conditions with the methionine source provided as the L-[methyl-^2^H_3_]-methionine isotope tracer. Following treatment up to 72 hours, genomic DNA was extracted, underwent an enzymatic nucleoside digestion, and the resulting free nucleosides were analyzed by LC-MS. From this tracing approach, the [methyl-^2^H_3_]-methionine produces [methyl-^2^H_3_]-SAM and when this isotope-labeled SAM is used as a substrate by DNMTs, this results in [methyl-^2^H_3_]-deoxycytidine on DNA (Fig. 2B). Using this LC-MS based approach, we successfully detected unmodified deoxycytidine and 5-methyldeoxycytidine, along with other unmodified nucleosides, from isolated DNA (Fig. S2A).

**Figure 2.**
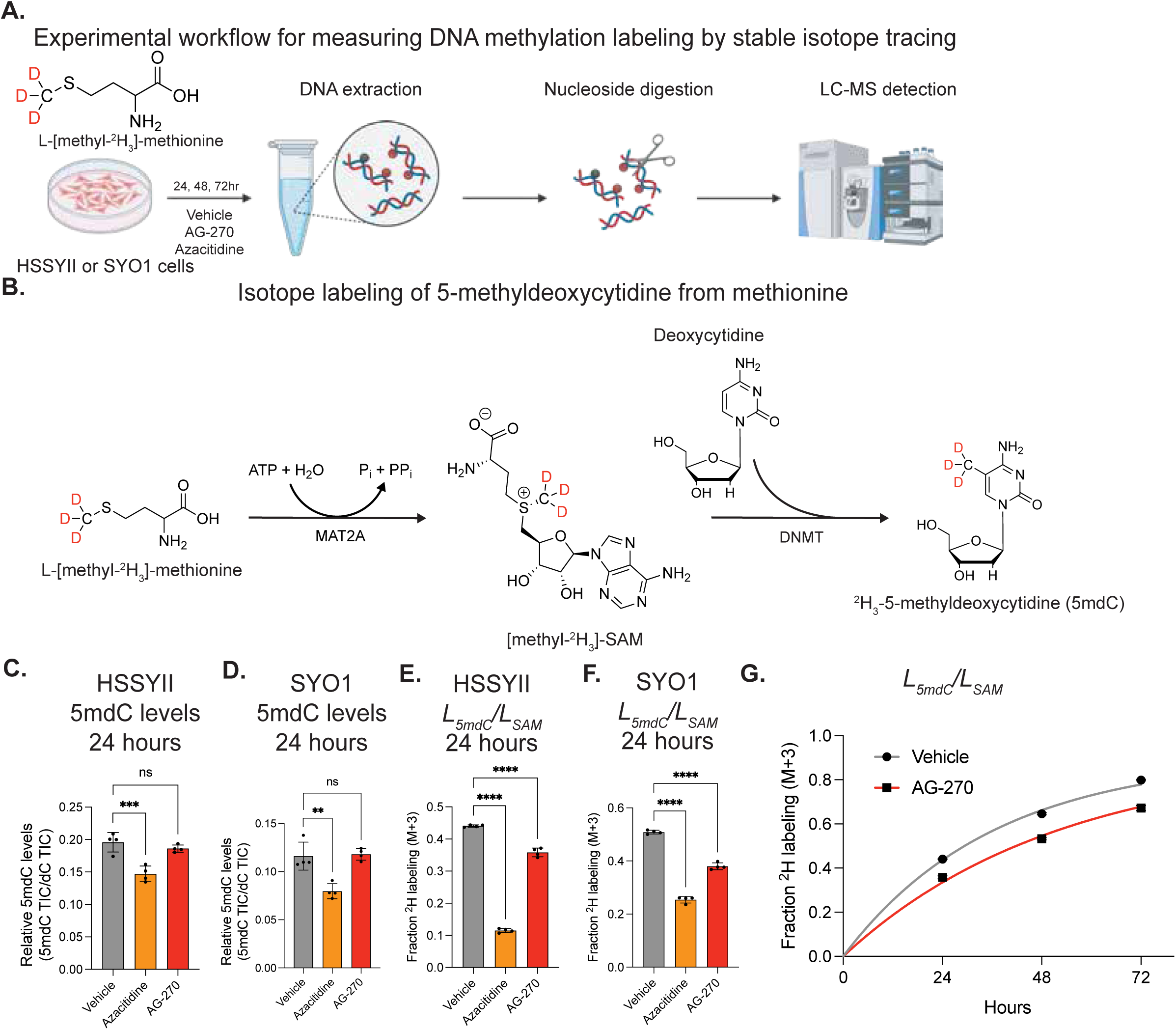
Stable isotope tracing shows MAT2A inhibition reduces turnover flux of 5-methyldeoxycytidine. (A) Experimental schematic of culturing cells with L-[methyl-^2^H_3_]-methionine with vehicle, azacitidine, or AG-270 treatment, isolating DNA, performing a nucleoside digestion, and analyzing nucleosides by LC-MS. (B) Schematic showing transfer of isotope labeled methyl group from methionine to SAM to 5-methyldeoxycytidine of DNA. (C-D) Levels of 5-methyldeoxycytidine measured by LC-MS after 24 hours of treatment with vehicle, 1 µM azacitidine, or 6 µM AG-270 in (C) HSSYII (n=4) and (D) SYO1 (n=4). (E-F) Fractional isotope labeling of 5-methyldeoxycytidine normalized to SAM labeling after 24 hours with tracer and vehicle, 1 µM azacitidine, or 6 µM AG-270 treatment in (E) HSSYII (n=4) and (F) SYO1 (n=4). (G) Fractional isotope labeling of 5-methyldeoxycytidine normalized to SAM labeling after 24-72 hours with tracer and vehicle or 6 µM AG-270 treatment in HSSYII (n=4). All data are reported as mean ± SD, p value by one-way ANOVA with Dunnett’s multiple comparison test and replicates (n) indicated. * denotes p < 0.05, ** denotes p < 0.01, *** denotes p < 0.001, **** denotes p < 0.0001. Data in (G) is fitted to a one-phase association exponential curve.

As expected, azacitidine treatment reduced total 5-methyldeoxycytidine levels within 24 hours (Fig. 2C-D). However, SAM depletion by AG-270 treatment caused no change in 5-methyldeoxycytidine levels for up to 72 hours (Fig. 2C-D, Fig S2B-C). Changes in metabolite concentrations, and in this case total DNA methylation levels, require a mismatch in synthesis and consumption fluxes(58). To measure the rate of DNA methylation, we quantified the labeling fraction of 5-methyldeoxycytidine of DNA from the L-[methyl-^2^H_3_]-methionine tracer. From this, we found that labeling of 5-methyldeoxycytidine, when normalized to the labeling percentage of SAM, was reduced following either DNMT or MAT2A inhibition (Fig. 2E-F, Fig. S2D-I). In fact, cells with a decreased SAM supply had a reduction in the rate of label incorporation into 5-methyldeoxycytidine over 72 hours (Fig. 2G). This suggests that depleting SAM levels caused a reduction in DNA methylation flux with a corresponding decrease in demethylation flux leading to slowed DNA methylation turnover while total methylation level remained constant.

### Reduced SAM levels induce a genomic redistribution of DNA methylation

To understand how this decrease in bulk DNA methylation turnover flux altered DNA methylation patterns genome-wide, we performed Enzymatic Methyl-sequencing (EM-seq) (Fig. 3A). Consistent with the previous LC-MS measurements, SAM depletion caused no change in the number of methylated CpG islands (Fig. S3A-B). As expected, inhibition of DNA methylation by azacitidine caused a dramatic and nearly universal loss of methylation at regions across the genome (Fig. 3B). Conversely, MAT2A inhibition resulted in an equal number of gained and lost methylated sites (Fig. 3C). When examining the genomic location of the regions with lost methylation, AG-270 treatment showed an enrichment for gene promoters compared to azacitidine (Fig. 3D). This suggests that the cellular response to reduced SAM supply is loss of methylation at certain genomic loci, while having a concurrent gain and placement of methyl groups at distinct genomic regions.

**Figure 3.**
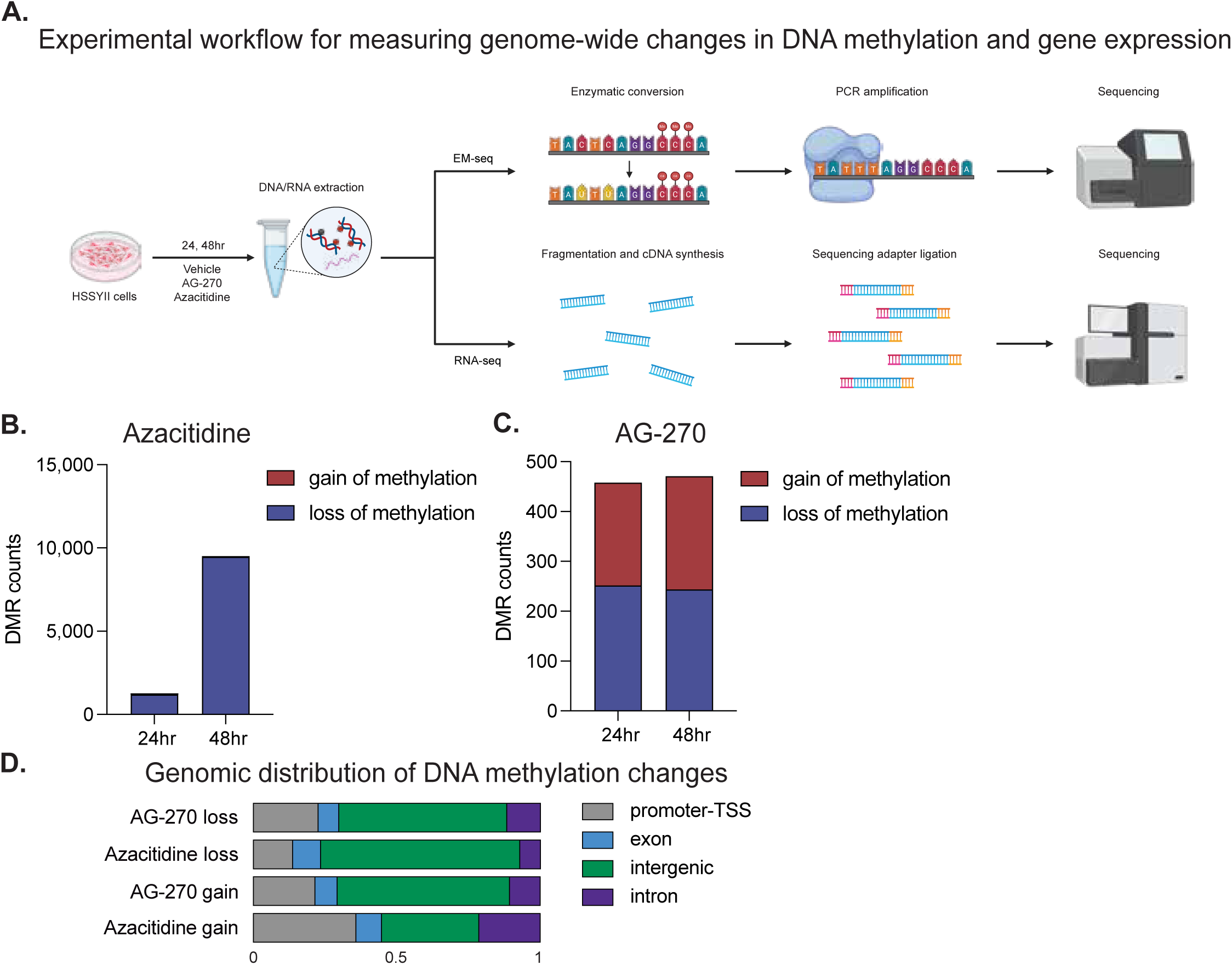
SAM depletion causes a genomic redistribution of DNA methylation. (A) Experimental schematic of treating cells with vehicle, azacitidine, or AG-270 followed by isolation of DNA for EM-seq analysis and mRNA for RNA-seq analysis. (B-C) Number of genomic regions with gain or loss of DNA methylation compared to vehicle following 24 and 48 hours treatment of HSSYII with (B) 1 µM azacitidine (n=3) and (C) 6 µM AG-270 (n=3). (D) Distribution of DNA methylation changes across genomic regions in HSSYII cells after 24 hours treatment of 1 µM azacitidine and 6 µM AG-270.

### SAM depletion causes transcriptional changes distinct from inhibition of DNA methylation

Since DNA methylation influences chromatin structure and accessibility, we performed transcriptomics analysis (RNA-seq) to understand how changes in SAM availability alter gene expression in HSSYII cells (Fig. 3A). MAT2A inhibition by AG-270 caused gene expression changes distinct from azacitidine after 24 and 48 hours of treatment (Fig. 4A, Fig S4A). Differential gene expression analysis compared to vehicle control showed both AG-270 and azacitidine treatment caused a greater number of genes to be upregulated than downregulated with limited overlap in the altered genes between treatment groups (Fig. 4B-C). Pathway enrichment analysis revealed that a primary result of SAM depletion was activation of transcriptional programs related to p53 activity (Fig. 4D-E). Despite the stark differences in DNA methylation patterning and global transcriptional profiles, both SAM depletion and azacitidine treatment converged to overcome the SS18-SSX-mediated block of differentiation to induce the cells into a more mesenchymal state, as indicated by the enrichment for genes associated with the Epithelial-Mesenchymal Transition (EMT) (Fig. 4F, Fig S4B)(52). Consistent with this mesenchymal activation leading to a reduction in cell proliferation, both treatments suppressed cell cycle progression and elevated apoptosis expression signatures (Fig. 4G-H, Fig S4C-D).

**Figure 4.**
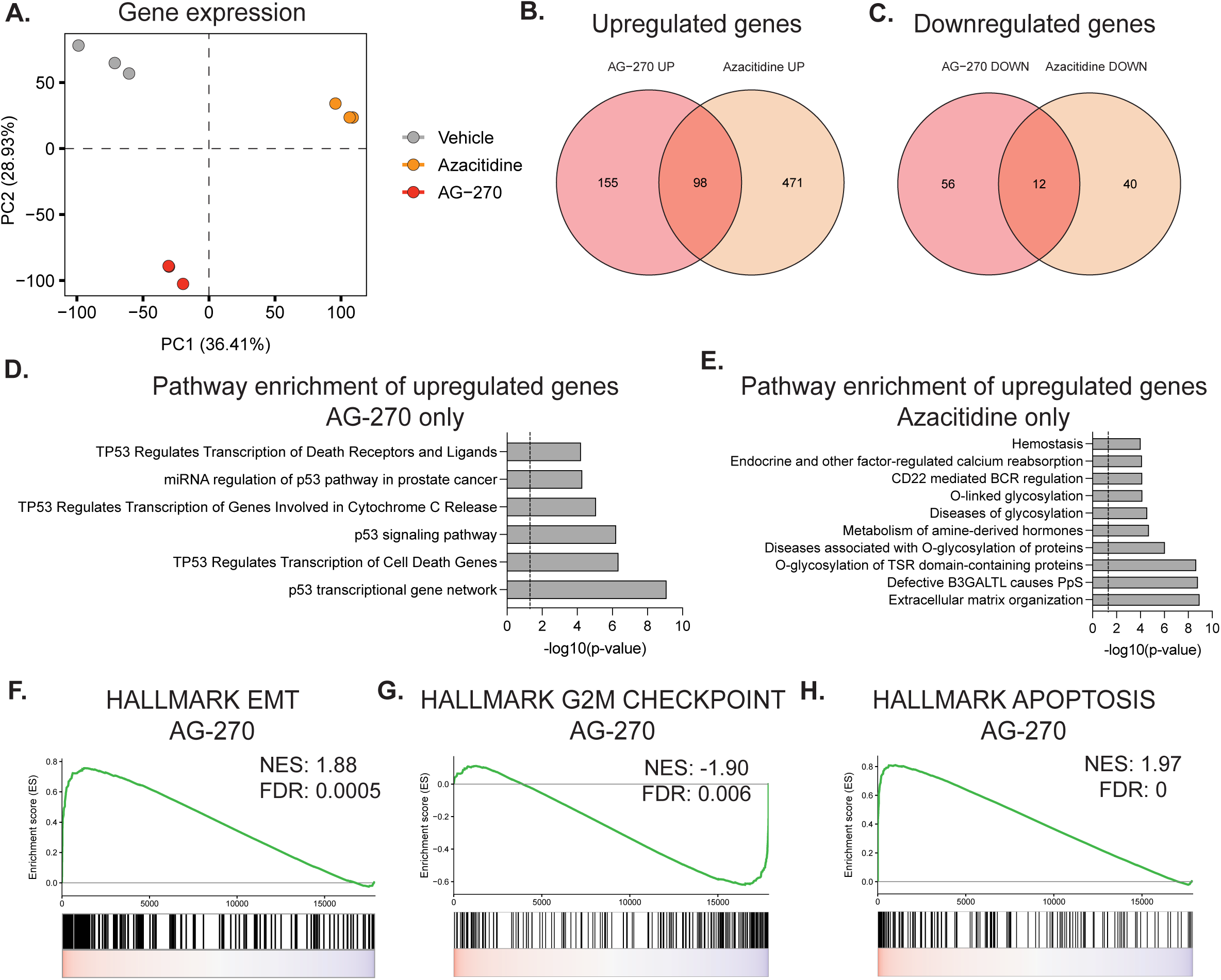
Inhibition of SAM synthesis induces transcriptional changes distinct from DNA methylation inhibition to induce differentiation in synovial sarcoma. (A) Principle component analysis (PCA) plot of gene expression in HSSYII after 48 hours treatment with vehicle, 1 µM azacitidine, or 6 µM AG-270 (n=3). (B) Overlap of differentially upregulated genes (log2FC > 1, adj p value < 0.05) in HSSYII treated for 48 hours with 1 µM azacitidine or 6 µM AG-270 compared to vehicle. (C) Overlap of differentially downregulated genes (log2FC < -1, adj p value < 0.05) in HSSYII treated for 48 hours with 1 µM azacitidine or 6 µM AG-270 compared to vehicle. (D) Gene ontology (GO) term enrichment analysis of the 155 genes upregulated only by 6 µM AG-270 treatment. (E) Gene ontology (GO) term enrichment analysis of the 471 genes upregulated only by 1 µM azacitidine treatment. (F-G) GSEA plots of HSSYII after 48 hours treatment with 6 µM AG-270.

### Transcriptional response to SAM depletion is independent of altered promoter methylation

In addition to DNA, methylation of multiple macromolecules, including histones and mRNA, combine to regulate transcriptional activity. To understand what gene expression changes caused by SAM depletion are directly associated with altered DNA methylation, we identified all genes with a change in methylation within their promoter region, defined as within 1 kb of the transcription start site. Azacitidine treatment caused a progressive disruption to promoter methylation with 36 genes at 24 hours and 5,284 genes at 48 hours with lost methylation (Fig. 5A-B). This loss in promoter methylation was associated with an increase in expression of these genes caused by azacitidine treatment (Fig. 5A-B). Conversely, the genes with altered promoter methylation caused by SAM depletion had no global change in expression (Fig. 5C-D). This data supports that the major driver of transcriptional changes in response to SAM depletion is through mechanisms beyond static level of DNA promoter methylation, including turnover rate and alterations in histone and mRNA methylation or nutrient sensing signaling programs(59).

**Figure 5.**
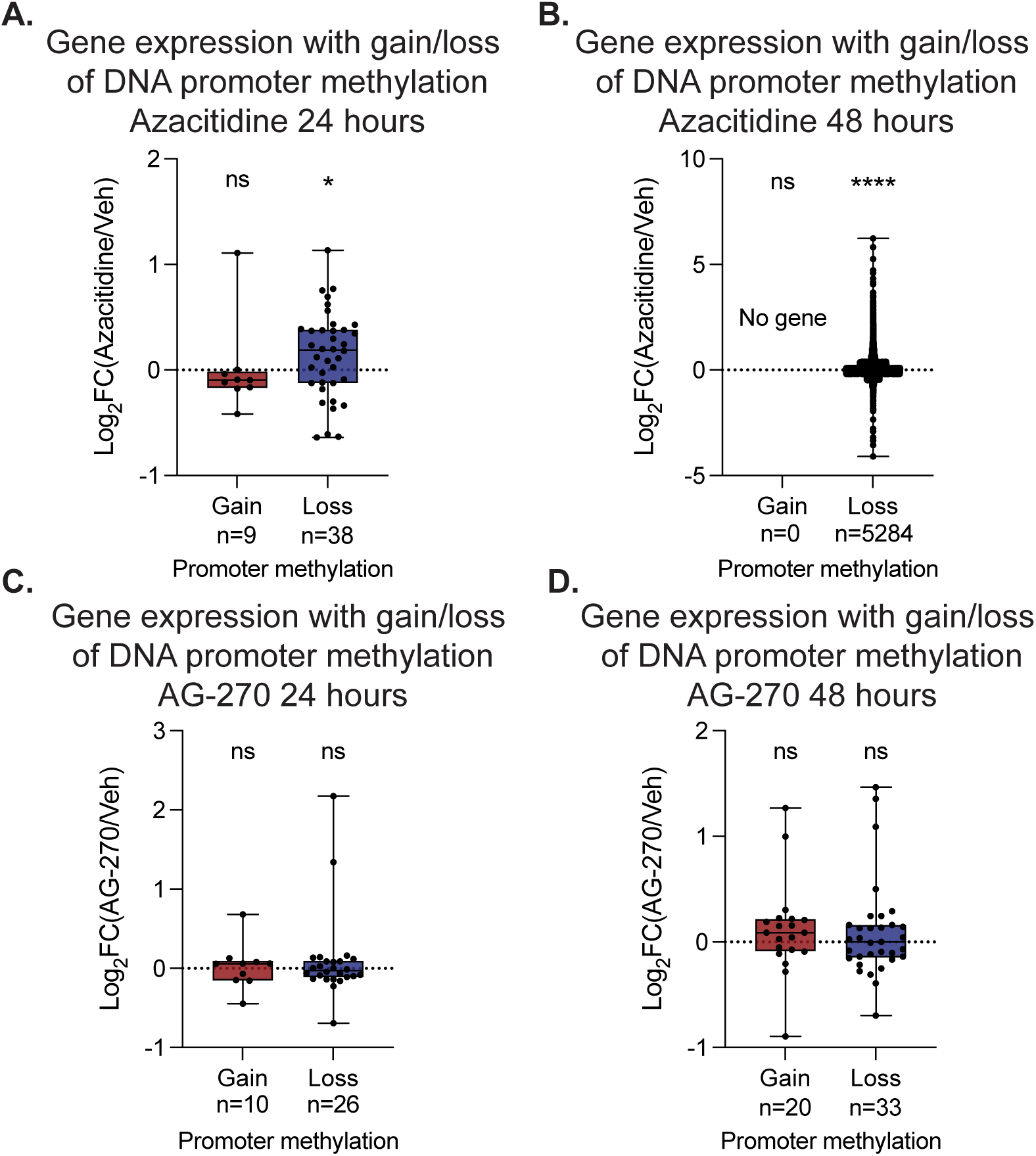
Altered gene expression caused by reduced SAM supply are independent of changes to DNA promoter methylation. (A) Gene expression change following gain or loss of DNA methylation at the gene promoter following 1 µM azacitidine treatment for 24 hours in HSSYII. (B) Gene expression change following gain or loss of DNA methylation at the gene promoter following 1 µM azacitidine treatment for 48 hours in HSSYII. (C) Gene expression change following gain or loss of DNA methylation at the gene promoter following 6 µM AG-270 treatment for 24 hours in HSSYII. (D) Gene expression change following gain or loss of DNA methylation at the gene promoter following 6 µM AG-270 treatment for 48 hours in HSSYII. For A-D, p value by one sample t and Wilcoxon test. ns denotes p > 0.05, * denotes p < 0.05, ** denotes p < 0.01, *** denotes p < 0.001, **** denotes p < 0.0001.

## DISCUSSION

Here we used stable isotope tracing to directly measure DNA methylation turnover fluxes and show that limiting SAM supply through MAT2A inhibition slows the rate of 5-methyldeoxycytidine turnover without changing its steady-state level. This kinetic effect, rather than a net loss of methylation, produced a genome-wide redistribution of DNA methylation with an equal number of losses and gains at distinct CpG islands. This was mechanistically and transcriptionally distinct from the bulk hypomethylation caused by azacitidine treatment. Disrupting SAM synthesis was sufficient to induce mesenchymal differentiation, arrest proliferation, and activate p53-associated transcriptional programs in synovial sarcoma cells, identifying SAM metabolism as a requirement for SS18-SSX-driven oncogenic growth.

The dogma that DNA methylation is a static process has been revised by work showing continuous turnover at specific genomic loci, with high-turnover sites disproportionately associated with active transcription(38, 41). Our data extend this concept by showing that the rate of turnover is under metabolic control and sensitive to the activity of the methionine cycle to supply SAM. Reducing intracellular SAM to low micromolar concentrations, in the range of the reported K_m_ values of DNMT1 and DNMT3A/B for SAM, slowed incorporation of labeled methyl groups into 5-methyldeoxycytosine without depleting bulk levels over the same timeframe. To prevent this depletion in bulk DNA methylation levels, demethylation rates must have decreased(60). This kinetic competition model predicts that different genomic loci have varied demand for SAM to support continuous re-methylation to counteract ongoing demethylation(61, 62). As a result, a reduction in SAM availability did not cause uniform hypomethylation, but drove a redistribution where sites with high demethylation activity lost methylation while other distinct loci gained methylation(61). This gain in methylation could presumably be due to lower competing demethylation activity or active regulation in response to nutrient sensing leading to increase DNMT activity at these genomic sites. This is a fundamentally different regulatory mechanism than genetic or pharmacological disruption of methyltransferase activity to block placement of new methyl groups, and argues that DNA methylation at both the bulk level and at the resolution of specific loci can be viewed as a flux-based system with tunable kinetics based on enzyme activity and substrate availability.

A central observation was that gene expression was independent of alterations in promoter methylation levels caused by SAM depletion. This indicates that the dominant driver of the transcriptional response to SAM restriction operates through a pathway other than static promoter DNA methylation. This is consistent with SAM depletion slowing global DNA methylation turnover rather inducing global DNA hypomethylation. Future work will need to identify the specific genomic sites with the greatest reduction in DNA methylation turnover, driving this global decrease in turnover, and determine the relationship with their transcriptional output. This is critical to consider when attempting to interrogate how B vitamin deficiencies and genetic alterations to one-carbon metabolism enzymes are ultimately driving transcriptional changes(29, 63). This work shows that likely DNA methylation turnover rates, rather than solely static DNA levels, caused by alterations in methionine cycle fluxes, are major contributors to how folate deficiency or MTHFR activity can epigenetically control gene expression(64).

Reduced SAM availability is also altering mRNA and histone tail methylation levels and dynamics, which combined with alterations in DNA methylation turnover, is contributing to changes in chromatin structure and accessibility(7, 65). In fact, reduced histone arginine methylation can elevate DNA damage, which may account for the p53 pathway activation signature caused by MAT2A inhibition(66). Future studies measuring changes in histone tail methylation and mRNA transcript methylation are required to fully understand the global control of SAM levels on regulating chromatin structure and gene expression.

An established hallmark of the SS18-SSX oncogenic fusion in synovial sarcoma is suppression of mesehnchymal differentiation(51). Genetic loss of the SS18-SSX fusion results in activation of mesenchymal differentiation genes in synovial sarcoma cell lines(50). Previous work with DNMT inhibitors, including azacitidine, show a pharmacological-based way to overcome this SS18-SSX-mediated blockage in differentiation(52). Our data show that limiting SAM supply also caused activation of mesenchymal genes, cell-cycle arrest, and elevated apoptotic signatures, yet through a mechanistically distinct route. Where azacitidine treatment caused a near-universal loss of methylation across the genome, MAT2A inhibition produced a balanced exchange of methylation gains and losses. The differentially expressed gene sets induced by each treatment showed limited overlap, indicating that SAM depletion does not simply produce an attenuated version of the DNMT-inhibition transcriptional program. Rather, phenotypic convergence appears to arise from largely non-overlapping transcriptional changes, suggesting that synovial sarcoma cells can be pushed toward a similar differentiated, non-proliferative state through targeting metabolic supply of SAM.

These findings establish cellular SAM supply as a regulator of DNA methylation turnover rate, distinct from its role in setting steady-state methylation level, and show that this kinetic control is sufficient to redistribute methylation genome-wide and disrupt oncogenic transcription in synovial sarcoma. This suggests that metabolic control of epigenetic methylation flux is a key mediator for how alterations in one-carbon metabolism can influence transcription with implications for B vitamin deficiency, inborn errors of metabolism, and cancer.

## ACKNOWLEDGMENTS

We are grateful to members of the McBride laboratory for insightful discussions, the members of the Kagan and Brunetti labs for assistance operating the SCIEX QTRAP LC-MS/MS instrument, and the members of the Metabolomics Shared Resource at Rutgers Cancer Institute for assistance with nucleoside detection by LC-MS.

## AUTHOR CONTRIBUTIONS

Y.L., M.S.S., M.E.J., M.E.D and K.K.N methodology; Y.L., M.S.S., M.E.J., L.K., L.B., X.S., and M.J.M. writing-review and editing; Y.L., M.E.J., and M.J.M. formal analysis, Y.L. and M.J.M. conceptualization; M.J.M. supervision; L.K., L.B., X.S., and M.J.M. resources; Y.L. and M.J.M. writing-original draft; M.J.M funding acquisition.

## DATA AND CODE AVAILABILITY

El-Maven (Elucidata) and AccuCor Isotope Natural Abundance Correction used for the analysis of LC-MS data is available on GitHub (https://github.com/lparsons/accucor). LC-MS metabolomics data will be deposited at the National Metabolomics Data Repository. Sequencing data from RNA-seq and EM-seq will be deposited to GEO.

## DECLARATION OF GENERATIVE AI IN SCIENTIFIC WRITING

During the preparation of this work, the authors used Heureka Bench (v4.4.0) in order to edit text for clarity. After using this tool/service, the authors reviewed and edited the content as needed and take full responsibility for the content of the publication.

## FIGURE LEGENDS

**Supplemental Figure 1.** (A) HSSYII synovial sarcoma cell line growth following vehicle, 3 or 6 µM AG-270, or 1 µM azacitidine treatment for 3 days and (B) 6 days (n=3). (C) SYO1 synovial sarcoma cell line growth following vehicle, 3 or 6 µM AG-270, or 1 µM azacitidine treatment for 3 days and (D) 6 days (n=3). (E) SAM total ion count quantified by LC-MS/MS in HSSYII after vehicle, 1 µM azacitidine, or 6 µM AG-270 treatment for 24 hours. (F) Intracellular SAM concentrations quantified by LC-MS/MS in HSSYII after vehicle or 6 µM AG-270 treatment for 24, (G) 48, and (H) 72 hours (n=4). All data are reported as mean ± SD, p value by one-way ANOVA with Dunnett’s multiple comparison test and replicates (n) indicated. * denotes p < 0.05, ** denotes p < 0.01, *** denotes p < 0.001, **** denotes p < 0.0001.

**Supplemental Figure 2.** (A) Nucleosides detected by LC-MS in HSSYII cells after 24 hours of treatment with vehicle (n=4). (B) Levels of 5-methyldeoxycytidine measured by LC-MS after 48 hours of treatment with vehicle, 1 µM azacitidine, or 6 µM AG-270 in HSSYII (n=4). (C) Levels of 5-methyldeoxycytidine measured by LC-MS after 72 hours of treatment with vehicle or 6 µM AG-270 in HSSYII (n=4). (D) Fractional isotope labeling of methionine and (E) SAM after 24 hours with tracer and vehicle, 1 µM azacitidine, or 6 µM AG-270 treatment in HSSYII (n=4). (F) Fractional isotope labeling of SAM normalized to methionine labeling after 24 hours with tracer and vehicle, 1 µM azacitidine, or 6 µM AG-270 treatment in HSSYII (n=4). (G) fractional isotope labeling of methionine and (H) SAM after 24 hours with tracer and vehicle, 1 µM azacitidine, or 6 µM AG-270 treatment in HSSYII (n=4). (I) Fractional isotope labeling of SAM normalized to methionine labeling after 24 hours with tracer and vehicle, 1 µM azacitidine, or 6 µM AG-270 treatment in HSSYII (n=4). All data are reported as mean ± SD, p value by one-way ANOVA with Dunnett’s multiple comparison test and replicates (n) indicated. * denotes p < 0.05, ** denotes p < 0.01, *** denotes p < 0.001, **** denotes p < 0.0001.

**Supplemental Figure 3.** (A) The percentage of methylated CpG islands in HSSYII cells after 24 hours and (B) 48 hours of vehicle, 1 µM azacitidine or 6 µM AG-270 (n=3). All data are reported as mean ± SD, p value by one-way ANOVA with Dunnett’s multiple comparison test and replicates (n) indicated. * denotes p < 0.05, ** denotes p < 0.01, *** denotes p < 0.001, **** denotes p < 0.0001.

**Supplemental Figure 4.** (A) Principle component analysis (PCA) plot of gene expression in HSSYII after 24 hours treatment with vehicle, 1 µM azacitidine, or 6 µM AG-270 (n=3). (B-D) GSEA plots of HSSYII after 48 hours treatment with 1 µM azacitidine.

