## Supplemental Figures for "S-adenosylmethionine supply controls DNA methylation turnover in synovial sarcoma"

**Supplemental Figure 1**

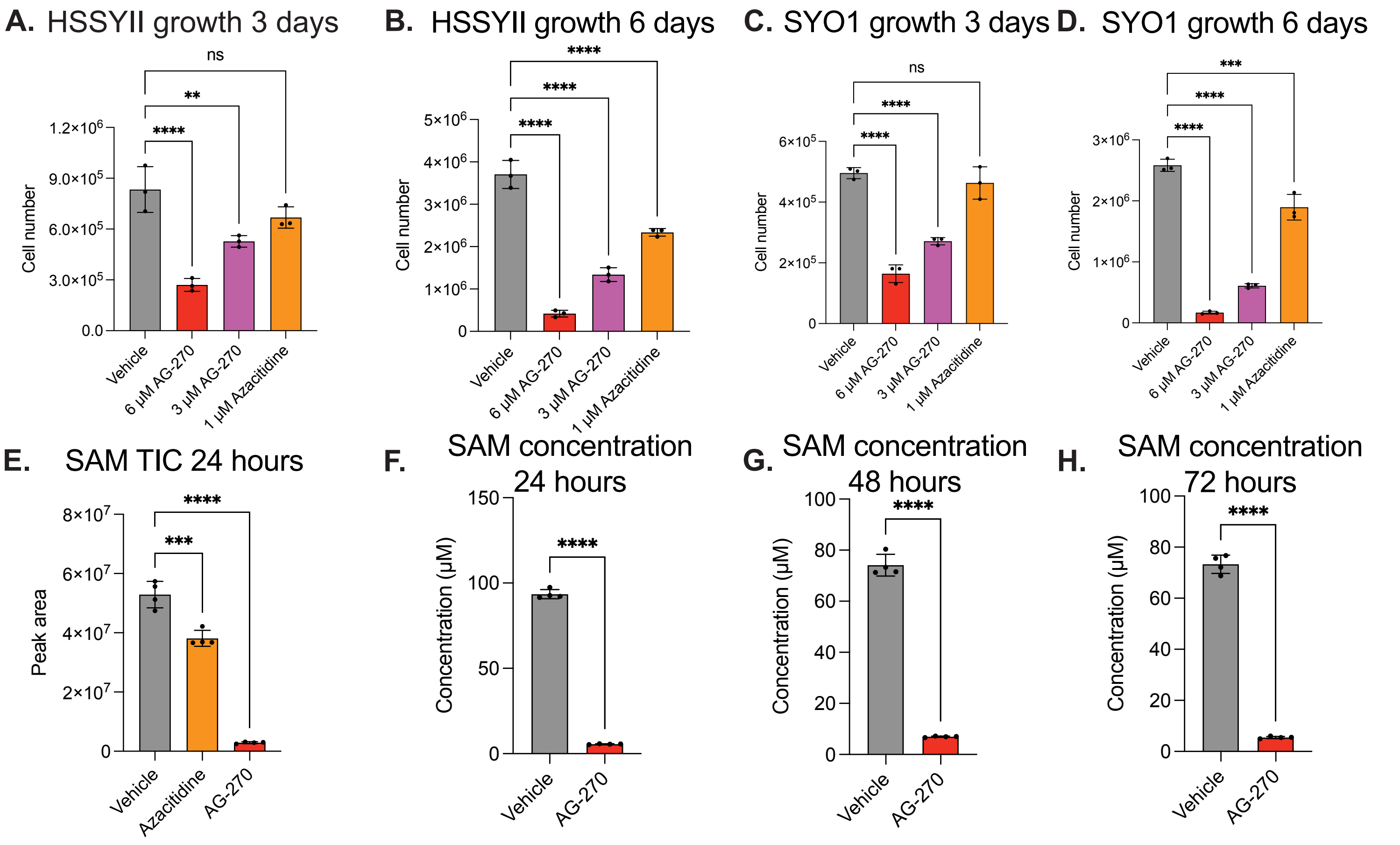


**Supplemental Figure 1.** (A) HSSYII synovial sarcoma cell line growth following vehicle, 3 or 6 µM AG-270, or 1 µM azacitidine treatment for 3 days and (B) 6 days (n=3). (C) SYO1 synovial sarcoma cell line growth following vehicle, 3 or 6 µM AG-270, or 1 µM azacitidine treatment for 3 days and (D) 6 days (n=3). (E) SAM total ion count quantified by LC-MS/MS in HSSYII after vehicle, 1 µM azacitidine, or 6 µM AG-270 treatment for 24 hours. (F) Intracellular SAM concentrations quantified by LC-MS/MS in HSSYII after vehicle or 6 µM AG-270 treatment for 24, (G) 48, and (H) 72 hours (n=4). All data are reported as mean ± SD, p value by one-way ANOVA with Dunnett's multiple comparison test and replicates (n) indicated. * denotes $p < 0.05$, ** denotes $p < 0.01$, *** denotes $p < 0.001$, **** denotes $p < 0.0001$.

### Supplemental Figure 2

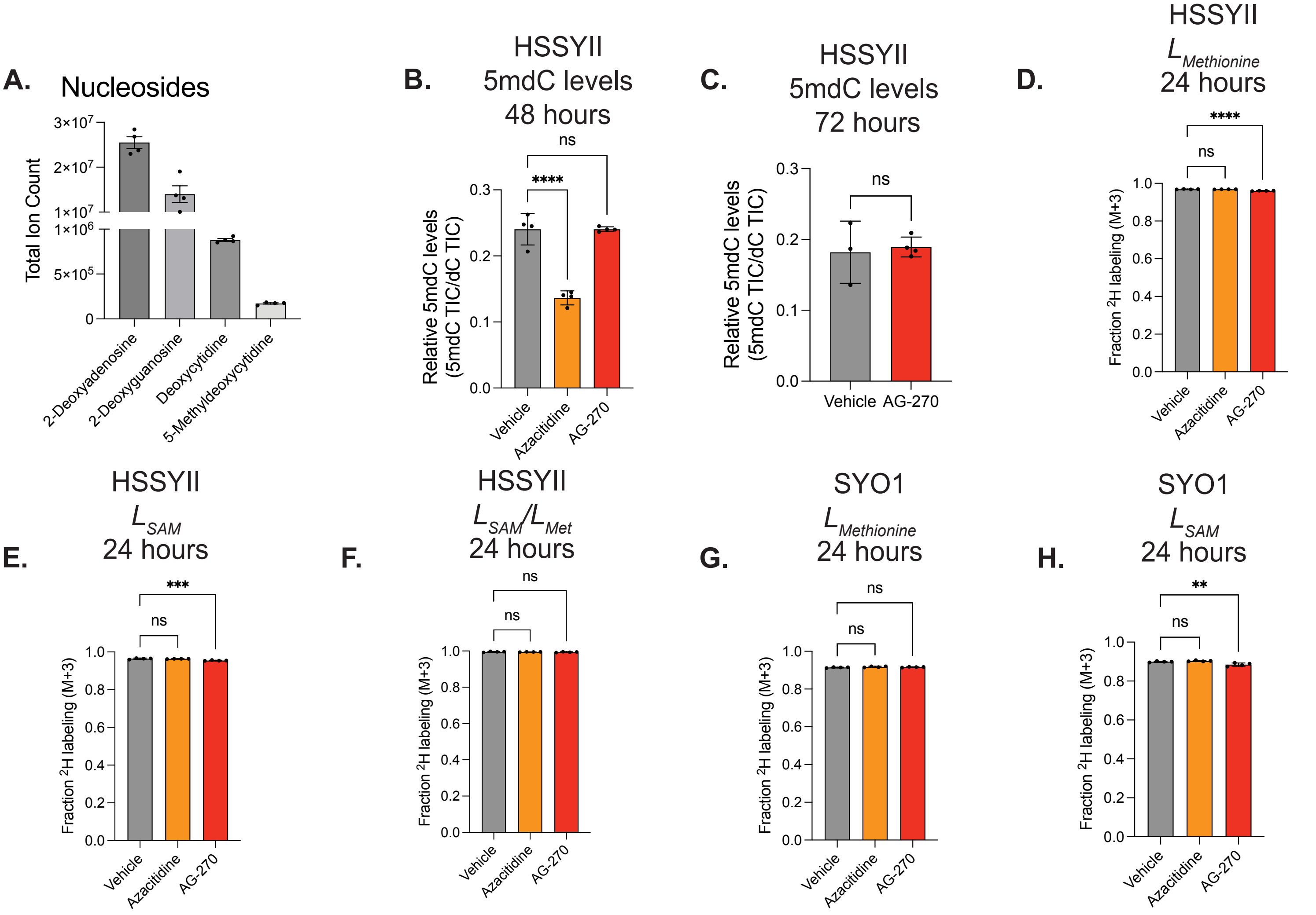


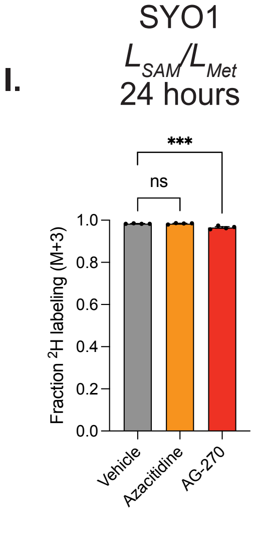


**Supplemental Figure 2.** (A) Nucleosides detected by LC-MS in HSSYII cells after 24 hours of treatment with vehicle (n=4). (B) Levels of 5-methyldeoxycytidine measured by LC-MS after 48 hours of treatment with vehicle, 1 µM azacitidine, or 6 µM AG-270 in HSSYII (n=4). (C) Levels of 5-methyldeoxycytidine measured by LC-MS after 72 hours of treatment with vehicle or 6 µM AG-270 in HSSYII (n=4). (D) Fractional isotope labeling of methionine and (E) SAM after 24 hours with tracer and vehicle, 1 µM azacitidine, or 6 µM AG-270 treatment in HSSYII (n=4). (F) Fractional isotope labeling of SAM normalized to methionine labeling after 24 hours with tracer and vehicle, 1 µM azacitidine, or 6 µM AG-270 treatment in HSSYII (n=4). (G) fractional isotope labeling of methionine and (H) SAM after 24 hours with tracer and vehicle, 1 µM azacitidine, or 6 µM AG-270 treatment in HSSYII (n=4). (I) Fractional isotope labeling of SAM normalized to methionine labeling after 24 hours with tracer and vehicle, 1 µM azacitidine, or 6 µM AG-270 treatment in HSSYII (n=4). All data are reported as mean ± SD, p value by one-way ANOVA with Dunnett's multiple comparison test and replicates (n) indicated. * denotes $p < 0.05$, ** denotes $p < 0.01$, *** denotes $p < 0.001$, **** denotes $p < 0.0001$.

**Supplemental Figure 3**

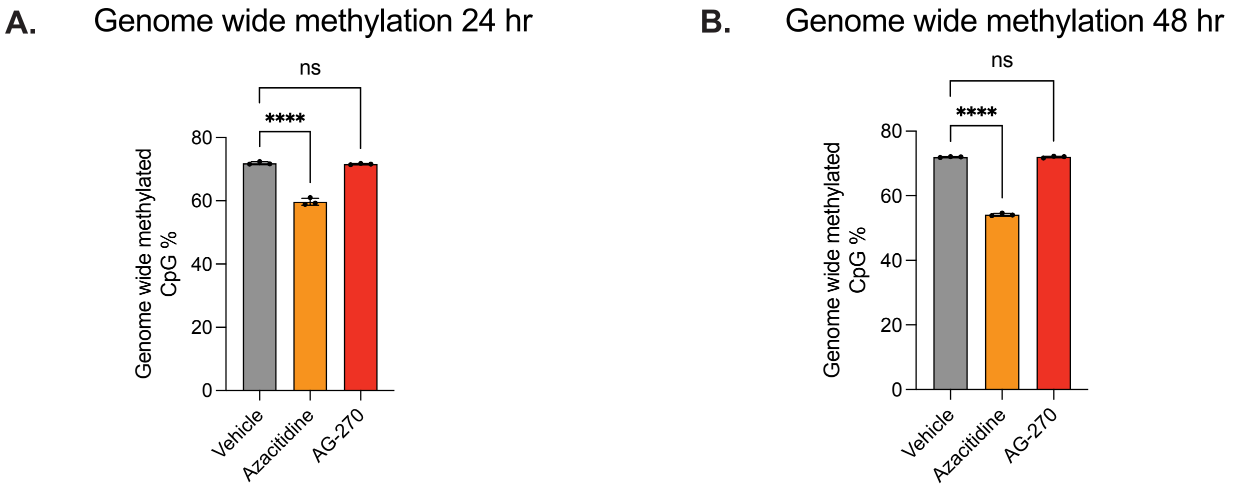


**Supplemental Figure 3.** (A) The percentage of methylated CpG islands in HSSYII cells after 24 hours and (B) 48 hours of vehicle, 1 μM azacitidine or 6 μM AG-270 (n=3). All data are reported as mean ± SD, p value by one-way ANOVA with Dunnett's multiple comparison test and replicates (n) indicated. * denotes $p < 0.05$, ** denotes $p < 0.01$, *** denotes $p < 0.001$, **** denotes $p < 0.0001$.

### Supplemental Figure 4

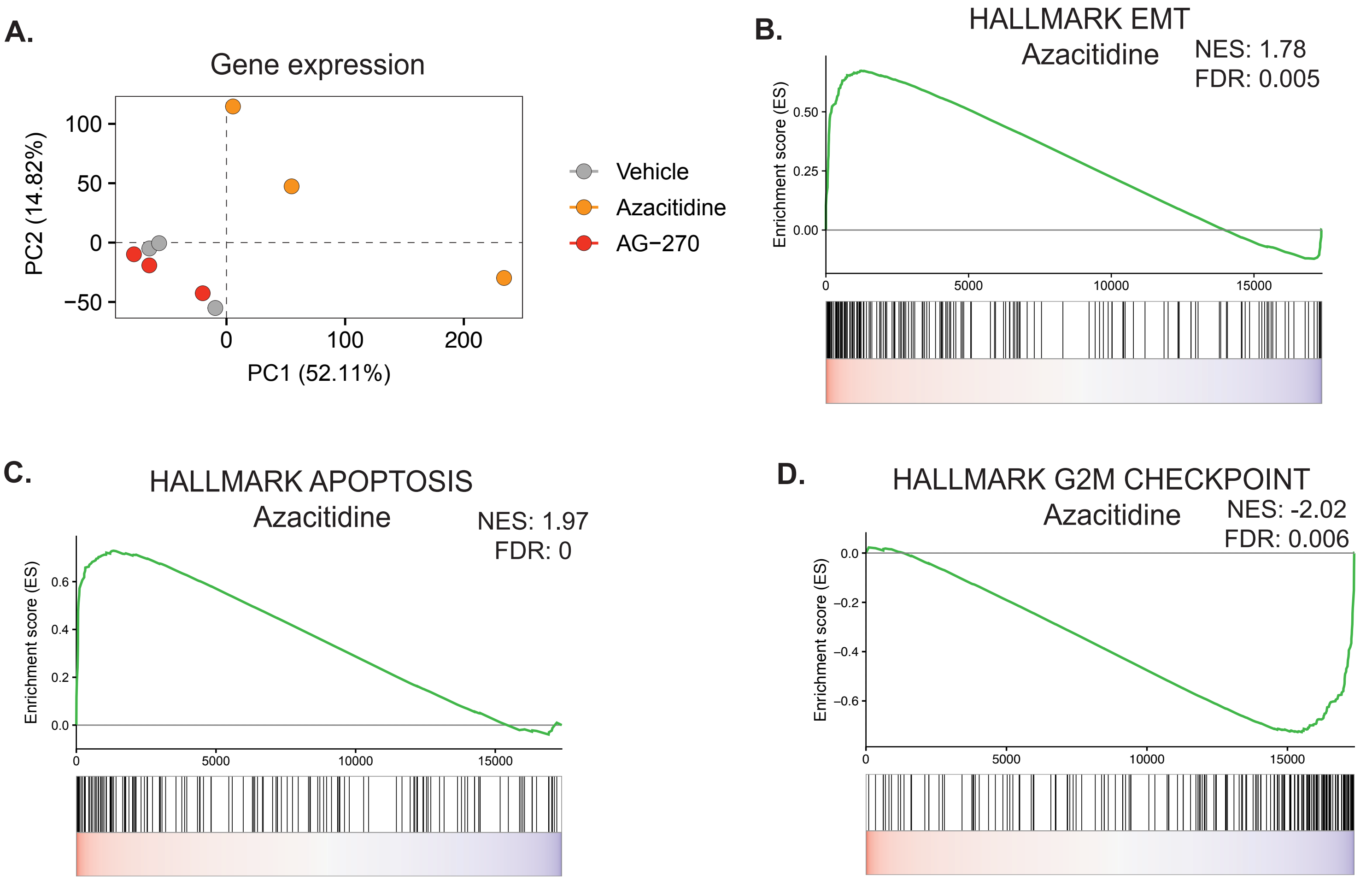


**Supplemental Figure 4.** (A) Principle component analysis (PCA) plot of gene expression in HSSYII after 24 hours treatment with vehicle, 1 μM azacitidine, or 6 μM AG-270 (n=3). (B-D) GSEA plots of HSSYII after 48 hours treatment with 1 μM azacitidine.
